# Cooperation and Stability for Complex Systems in Resource-Limited Environments

**DOI:** 10.1101/514018

**Authors:** Stacey Butler, James O’Dwyer

**Affiliations:** Department of Mathematics, University of Illinois, Urbana, IL 61801, USA; Department of Plant Biology, University of Illinois, Urbana, IL 61801, USA

## Abstract

Resource-limited complex systems are ubiquitous in the natural world, as is the potential for instability in such systems. Classic models of interacting species have provided a basis for our understanding of stability in these systems, and suggest that stable coexistence requires weak, rare, and asymmetric interactions. But missing from these models is an explicit understanding of how resource exchange and resource limitation can drive or prevent instability. Here we show that systems based on general rules for the consumption and exchange of resources are guaranteed to be stable when exchange of resources is reciprocated by each pair of partners. These cooperative, mutualistic interactions can be arbitrarily strong and yet not disrupt stability. More general modes of exchange will lead to instability when supply rates are low, but when resource supply from outside the system is sufficiently high, arbitrary exchange is consistent with a stable equilibrium.

## 1 Introduction

The general properties of large, interacting ecological systems have often been modeled using pairwise interactions between species, where changes in the population size of one species directly affect the growth rate of another (*33, 53*). While the strength of any given interaction is challenging to infer from empirical community data (*35, 45, 52*), simple null models have been used to gain general insights into equilibria and stability in these systems (*18, 37*). These results have shown that equilibria will tend to become unstable to small perturbations once either the number or strength of interactions passes a certain threshold (*2*), and that asymmetric mutualistic interactions enhance stability (*6*). Beyond strictly ecological applications, these insights have been leveraged in other complex biological and social networks comprising multiple, heterogenous component parts (*5, 14, 23, 46*). For example, while nodes in an ecological network are species and links typically represent interactions, in economic systems nodes have been interpreted as financial entities and links in terms of transactions between them (*38*). It is therefore fair to say that these stability results have deeply informed our understanding of interacting complex systems in general.

However, missing from this approach are the explicit mechanisms underlying interactions between individuals. Consumer-resource models meet this need by explicitly considering the consumption and preferences of individuals for resources (*1, 10, 11, 34, 44, 49–51*), and can lead to dynamics that differ from models based on direct species interactions (*39, 52*). Recent work has extended these analyses to large, open systems with extensive exchange of resources (*8,15,16,20,21,36*), and incorporating resource exchange as the mechanism underlying mutualistic interactions already leads to contradictions with the classical analyses using pairwise interactions. For example, if each species in a community specializes on a single resource, then local stability is guaranteed when each pair of species exchanges resources symmetrically (*8*), independent of how strong those interactions are. This counters both the idea that there is a limit on the strength and prevalence of interactions, and the result that asymmetry enhances stability in the case of mutualistic interactions, leveraged in both biological (*6*) and social systems (*38*).

These earlier results establish the importance of incorporating explicit resource exchange even when tackling basic questions related to equilibria and stability, but were limited in their scope to specific modes of consumption. For example (*8*) focused on consumers exchanging resources, while each specialized on consuming one particular resource. But in moving from models of direct species interactions to more mechanistic models of competition and mutualism, there is a tradeoff. Instead of describing an interaction with a single number, we now have a realm of different mechanisms to choose from, and any potentially general conclusions must be robust to different modes of consumption and exchange. Here, we begin to address the question of how robust are these earlier results, by allowing for relatively general forms for consumption and organism growth rates as a function of available resources. We use these models to address two open questions related to resource exchange in complex ecological communities. First, we show that the connection between reciprocity and stability is general, extending to more realistic models of consumption where each species’ growth is colimited by multiple, essential resources. We find that exact reciprocity is again sufficient for ecological stability, regardless of the strength or prevalence of exchange, and independent of the precise form of consumption. This generalization over previous results provides significant and essential support for the wider relevance of the reciprocity-stability connection.

Second, we consider the effects of environmental context on stability in terms of resource supply rates. This question has been raised in systems with resource subsidies from outside (*32, 41, 42*), but has not been addressed for large, complex communities exchanging resources arbitrarily. It has been unclear how to model the effect of a changing environment on interaction strengths in models of direct interactions (*9,56*), and so here, we consider the impact of the resource inflow rates on stability for communities that consume and exchange resources. We focus initially on the case of communities where consumers specialize on a single resource, and prove that in the limit of scarce resources any departure from a strict set of criteria for exchange will tend to destabilize the system. These criteria include reciprocity, alongside other limited modes of exchange. Conversely, for sufficiently large resource inflows, any mode of exchange can have a stable positive equilibrium. We subsequently generalize this proof to arbitrary modes of consumption, and show numerical examples for specific cases. These illustrate that when generic modes of consumption are combined with generic modes of exchange, the result can be compatible with a stable equilibrium, as long as resource inflows are sufficiently large, and suggest that in real ecological systems there will be important interplays between resource exchange, ecological stability, and environmental context.

## 2 Model of Consumption and Exchange

Our model of consumption and exchange is based on a set of *m* abiotic resources with densities *R*_*α*_, and *m* species with population sizes *N*_*i*_. We first give three classic models of consumption processes, in order to build up the intuition for what a reasonable consumption process look like. We’ll then prove a set of results that apply to each of these three examples, and also extend this to more general consumption processes that incorporate these three as special cases.

### 2.1 Specialism

In our first example, each species specializes on a single resource, and cannot grow without the resource being available. Exchange of resources is incorporated by allowing each species *i* to recycle a given proportion of its biomass back into the common pool, at a rate *P*_*αi*_ for resource *α*, while resource *α* is supplied to the system from outside at a specific rate *ρ*_*α*_ and removed (due to degredation or outflow) at rate *η*_*α*_. Species undergo mortality (with biomass not recycled into usable resources) at rates *µ*_*i*_.

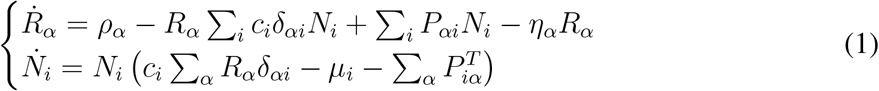

Here, *c*_*i*_ is the specific rate of consumption of species *i*, and the Kronecker delta symbol *δ*_*iα*_ indicates that consumption only occurs when *α* is equal to *i*. Note that the rates, {*P*_*αi*_} form a matrix, *P*, and so 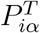 represents the row *i*, column *α* entry in the transpose of *P*.

These equations have equilibrium solutions

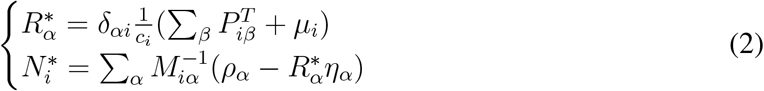

where 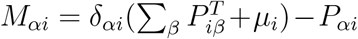. Not all combinations of *ρ*_*α*_ and *µ*_*i*_ will lead to positive (so-called feasible) equilibria. While the equilibrium solutions for 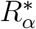 are always positive in this model, 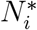 is not guaranteed to be positive. However, for a given set of 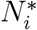, we may find {*ρ*_*α*_} and {*η*_*α*_} such that 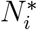 is a solution.

### 2.2 Substitutable Resources

In our second example, each species can consume multiple distinct resources, with uptake rates determined by *C*_*αi*_, which could also be thought of as the preference of species *i* for resource *α*. A given consumer still cannot grow without availability of the right resources, but can now have a wider range of options. Exchange of resources is again incorporated by allowing each species *i* to recycle a given proportion of its biomass back into the common pool, at a rate *P*_*αi*_ for resource *α*, with rates *ρ*_*α*_, *η*_*α*_ and *µ*_*i*_ defined as before.

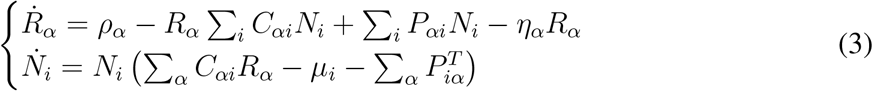

These equations have equilibrium solutions

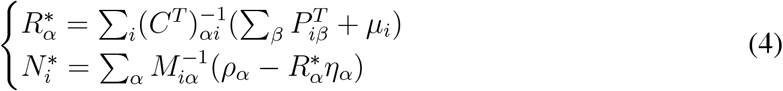

Where 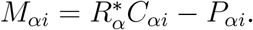.

Although in the model, feasibility is not guaranteed, for any given set of positive 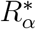, there exists a set of *µ*_*i*_ such that 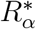 is a solution and for a given set of 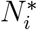, we may find {*ρ*_*α*_} and {*η*_*α*_} such that 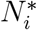 is a solution.

### 2.3 Colimitation by Multiple Resources

In this system, each species is colimited by a distinct set of resources, so that species *i* requires precisely *K*_*αi*_ units of resource *α* to grow new biomass. We model this consumption process using multiplicative colimitation (*13, 24, 39, 43*), though we also note that it would also be reasonable to consider other models of colimitation, for example Liebig’s law (*12, 19, 49, 54*), where each species has a single limiting resource at any given time. Each species *i* has a resource conversion rate of *r*_*i*_. Exchange of resources is as usual incorporated by allowing each species *i* to recycle a given proportion of its biomass back into the common pool, at a rate *P*_*αi*_ for resource *α*, with *ρ*_*α*_, *η*_*α*_ and *µ*_*i*_ retaining their previous definitions.

The resulting dynamics are described by the following ordinary differential equations:

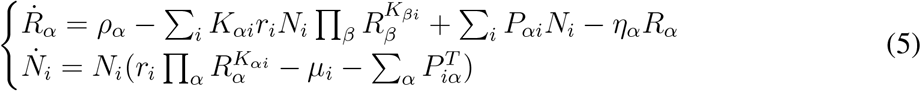

These equations have equilibrium solutions where consumption and production balance supply and mortality:

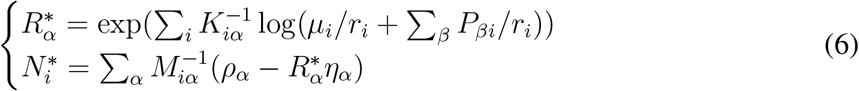

where the matrix *M* is given by *M*_*αi*_ = *K*_*αi*_(*µ*_*i*_ + Σ_*β*_ *P*_*βi*_) − *P*_*αi*_. Clearly not all combinations of *ρ*, *µ*, *r*_*i*_, *K* and *P* will lead to positive (so-called feasible) solutions. On the other hand, all positive values 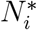 are possible for some choice of resource inflow *ρ*_*α*_. Conversely, not all resource concentrations are possible—the recycling of resources into the common pool typically places lower bounds on equilibrium resource concentrations for a given *K* and *P*, as discussed in our Appendices.

### 2.4 General Model of Consumption and Exchange

This variety in consumption processes simply reflects that when competition is modeled mechanistically, via the uptake and conversion of resources into biomass, there are then many distinct ways for this mechanism to occur. This contrasts with modeling competition implicitly, via direct species interactions (e.g. the competitive Lotka-Volterra equations), and means that while we may gain new insights by incorporating more explicit mechanism, we also have to deal with a potentially broader array of models. This in turn means that if we want to derive general results, for example about the stability of equilibria, we may need to make sure that these results hold for this broad range of models, and not just one special case.

With this in mind, we now define a more general model of consumption, incorporating as special cases the three examples above, but allowing for combinations of processes (e.g. the possibility that a consumer may generate biomass through more than one combination of essential resources—i.e. by one of multiple metabolic pathways). This model takes the form:

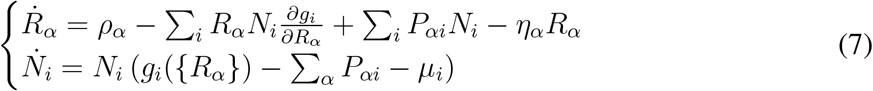

The freedom in choosing the consumption function, *g*_*i*_({*R*_*α*_}), as a function of resource concentrations allows us to straightforwardly recover any of the three earlier examples, in addition to a broad range of generalizations. Third, the form of the depletion of resources as a result of consumption generically takes the form 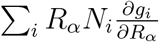 because we assume that the dependence of *g*_*i*_ on *R*_*α*_ captures the effective number of units of resource *α* required to build one new consumer, as in the case of multiplicative colimitation. We note that while covering a large set of plausible forms of consumption, and hence we describe it as general, this model is far from completely general. For example, a Michaelis-Menten form for resource uptake and depletion would not fall into this category; neither would the functional forms found in the models studied in (*27, 51*), where a species may consume more than one resource, but their growth rate is determined by one limiting resource (i.e., Liebig’s Law). In essence, what we generalize to here are cases where uptake is limited by the rate of *finding* all necessary resources moving diffusively in a given environment, so that it is not species’ intrinsic ability to metabolize nutrients that limits their growth rates. In real systems where resources are low, perhaps in oligotrophic environments, then we expect our approximation to work well. Finally, we put a restriction on the functional form of the consumption process, by requiring that the matrix with components

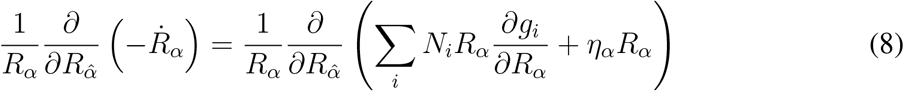

is a positive semi-definite matrix. More explicitly, this condition imposes that if we inflate or deflate resource concentrations by allowing *R*_*α*_ → *R*_*α*_(1 + *ϵ*_*α*_) for an arbitrary vector of small quantities *ϵ*_*α*_, then

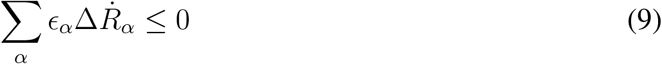

where 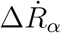 is the small change in growth rates 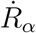 caused by the small change in resource concentrations *ϵ*_*α*_*R*_*α*_. As an example, if all resources are inflated by a the same factor, so that *ϵ*_*α*_ = *ϵ*, then this condition tells us that the sum of all depletion rates must increase. In essence, this condition imposes that resource depletion can’t decrease if resource concentrations increase.

### 2.5 Quantifying Species Interaction Strengths

In models where interactions are not mediated by explicit resources, the rate or strength of a species interaction is a part of the definition of the model. That’s not the case here, and so if we want to state or discuss the properties of these models and how they depend on species interactions, we first need the right definition of interaction strength. In earlier work on exchange among specialist consumers, it was natural to define the elements of the exchange matrix *P* as the strength of mutualism between each pair of species, because there was a one-to-one relationship between resource use and consumer identity. i.e. the entries of the exchange matrix *P* already gave us a natural definition of mutualism. However, in all of our more general cases, from substitutable resources through to the general consumption process encoded by Eq. (7), the ‘strength’ of mutualism is more subtle.

Therefore to make progress we now derive a metric to measure the strength of mutualism due to exchange between each pair of species. To simplify notation, we will in places use matrix-vector notation. Specifically, we define 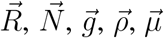 and 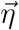 to represent vectors of the values of *R*_*α*_, *N*_*i*_, *g*_*i*_, *ρ*_*α*_, *µ*_*i*_ and *η*_*α*_ respectively. We define *R*_*d*_, *N*_*d*_ and *G*_*d*_ to be matrices formed by placing 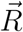, 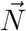 or 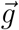 along the diagonal with zeros on the off-diagonal. We also define *Dg* by 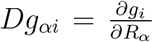 and we let 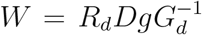. First, we interpret the matrix, *W*. Its components look like

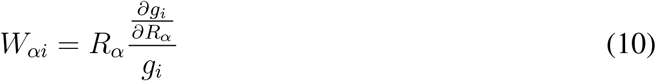

So that *W*_*αi*_ represents the current relative effect of resource *α* on species *i*’s growth rate multiplied by the current resource density. We assume that 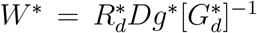, the matrix *W* evaluated at equilibrium, is of full rank, and hence invertible. In essence, this is an assumption that each species is distinct in terms of its phenotype. We can therefore define a matrix

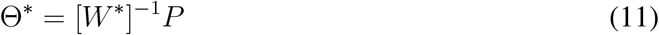

where *P* is the matrix describing the production of resources by each species. We note that this definition of interaction strength is reminiscent of earlier work characterizing empirical interactions in terms of resource flows (*22*).

We can interpret the components of Θ^*\**^ by considering their relationship to the components of *W*^*^ and *P*:

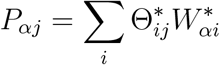

so that

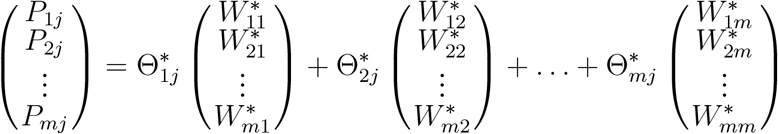

which can be written concisely as

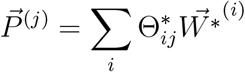

where 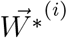 is a vector describing the resource requirements of species *i* (i.e. the *i*th column of the matrix *W*^*^) and 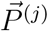 describes the production of resources by species *j*. It can be seen that the production of resources by *j* is a sum of components, each proportional to the resource requirements of another species, *i*, with the constant of proportionality equal to 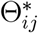.Hence, for all 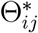 positive, we interpret 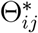 biologically as the strength of exchange from species *j* to species *i*. We note though that our sufficient condition for stability will require that all 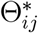 are non-negative, and so we restrict our analysis to systems with non-negative Θ^*\**^.

#### 2.5.1 Non-negative Mutualistic Interaction Strengths

We have shown that the production of resources by species *j* is a sum of components, each proportional to the resource requirements of another species, *i*, with the constant of proportionality equal to 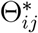. Hence, for all 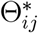 positive, we interpret 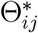 biologically as the strength of exchange from species *j* to species *i*. Our sufficient condition for stability will also require that all 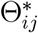 are non-negative (although we note that this is technically not a necessary condition), and so we consider it reasonable in this context to restrict our analyses to systems with non-negative Θ^*\**^. We also note that several of our examples also ensure that Θ is positive even away from equilibria.

## 3 Results

### 3.1 Reciprocity Implies Stability

Our model with generalized consumption, restated using matrix-vector notation is

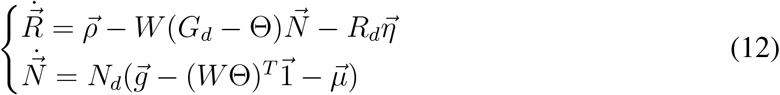

In our Appendices we prove that the equilibrium solution to (12) will be stable to small perturbations when the following conditions are met:

1. *Dg*^*^ is nonsingular
2. 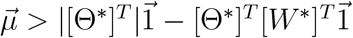
3. 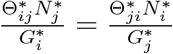 for all *i* and *j*

The first two conditions set bounds on species consumption and production. The first condition states that species’ responses to changes in resource densities should be sufficiently different. The second condition ensures that the eigenvalues of 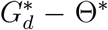 will be positive. This can be interpreted as the total strength of a species’ production/exchange being bounded by its consumption.

The third condition defines reciprocity. This is a condition on the structure of the exchange network. An exchange between two species, *i* and *j*, is reciprocal if species *j* produces as much of what species *i* needs relative to its growth rate, as species *i* does for species *j*. If this is true for all pairs *i* and *j*, then (when combined with the other conditions) the whole community will stably coexist. Note that although reciprocity is not a necessary condition for stability, it is not superfluous. Figures 1–3 show that in a range of cases where reciprocity is absent, there are transitions from stability to instablility as resource inflow is decreased. This suggests that generic positive interactions will not guarantee stability for all resource inflows, in contrast to the case of reciprocal positive interactions.

**Figure 1:**
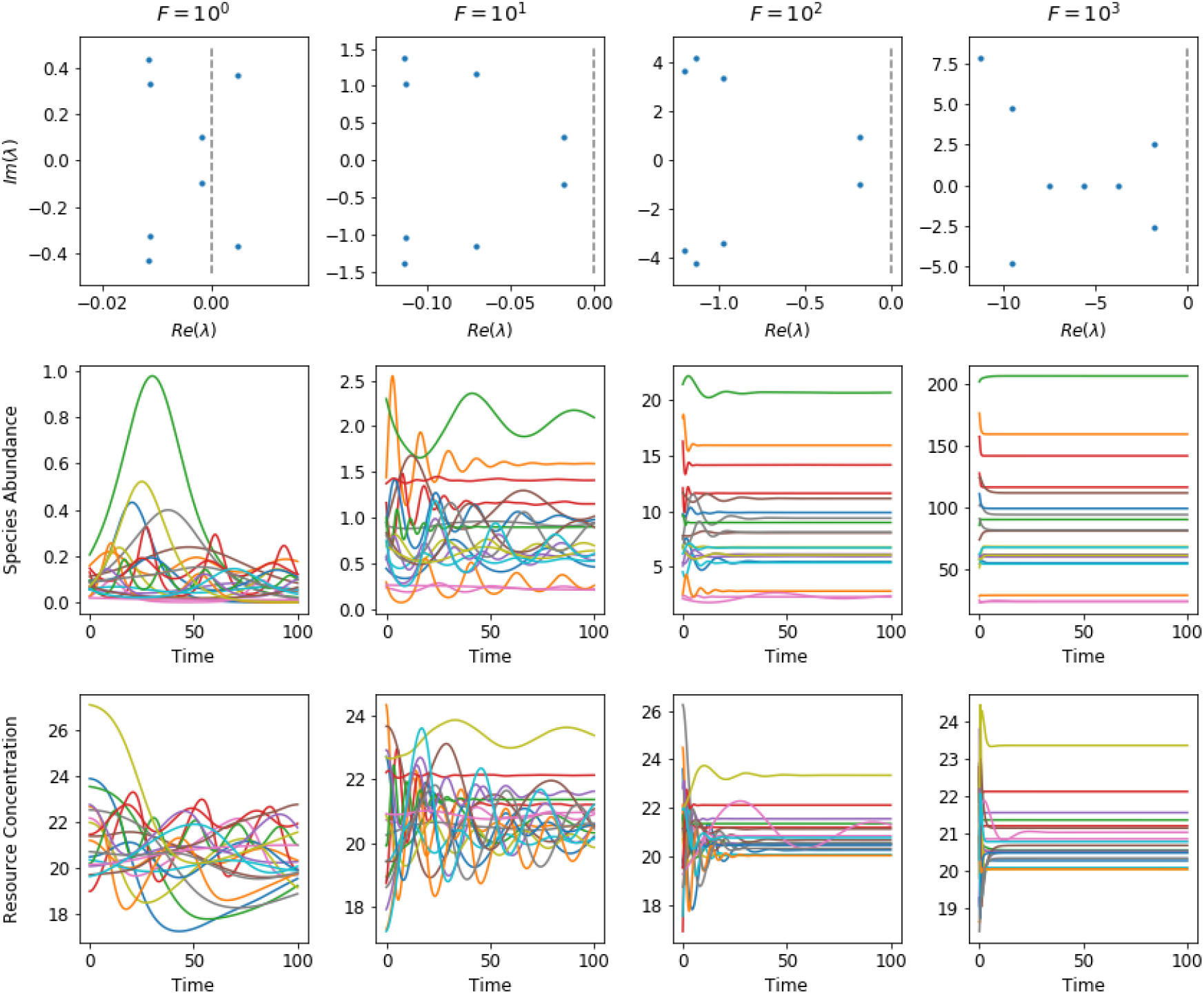
Resource Supply and the Onset of Instability for Specialist Consumers. This example shows four communities of 20 consumer species, each specializing on one of 20 distinct resources, with consumption rates drawn from the absolute values of a normal distribution with mean zero and standard deviation one. Inflow and mortality rates are chosen so that equilibrium concentrations and abundances are also randomly drawn from the absolute values of a normal distribution, and the exchange matrix Θ has a connectance =0.2, with non-zero entries drawn from the absolute values of a Gaussian (also with mean zero and sd one). From left to right we show the effect of changing resource supply on the dynamics of this community near equilibrium. Resource supply rates, 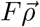, are increased by a factor of ten from each panel to the next, reflected in the increase of consumer abundances by the same factor. This shift demonstrates that reducing resource supplies alone can change a stable community equilibrium into an unstable equilibrium, mediated by the corresponding change in consumer abundances. The top panels illustrate the same phenomenon by focusing on the eigenvalue spectrum of the Jacobian near the equilibrium in each case. As resource supply is increased, the spectrum loses eigenvalues with positive real part.

**Figure 2:**
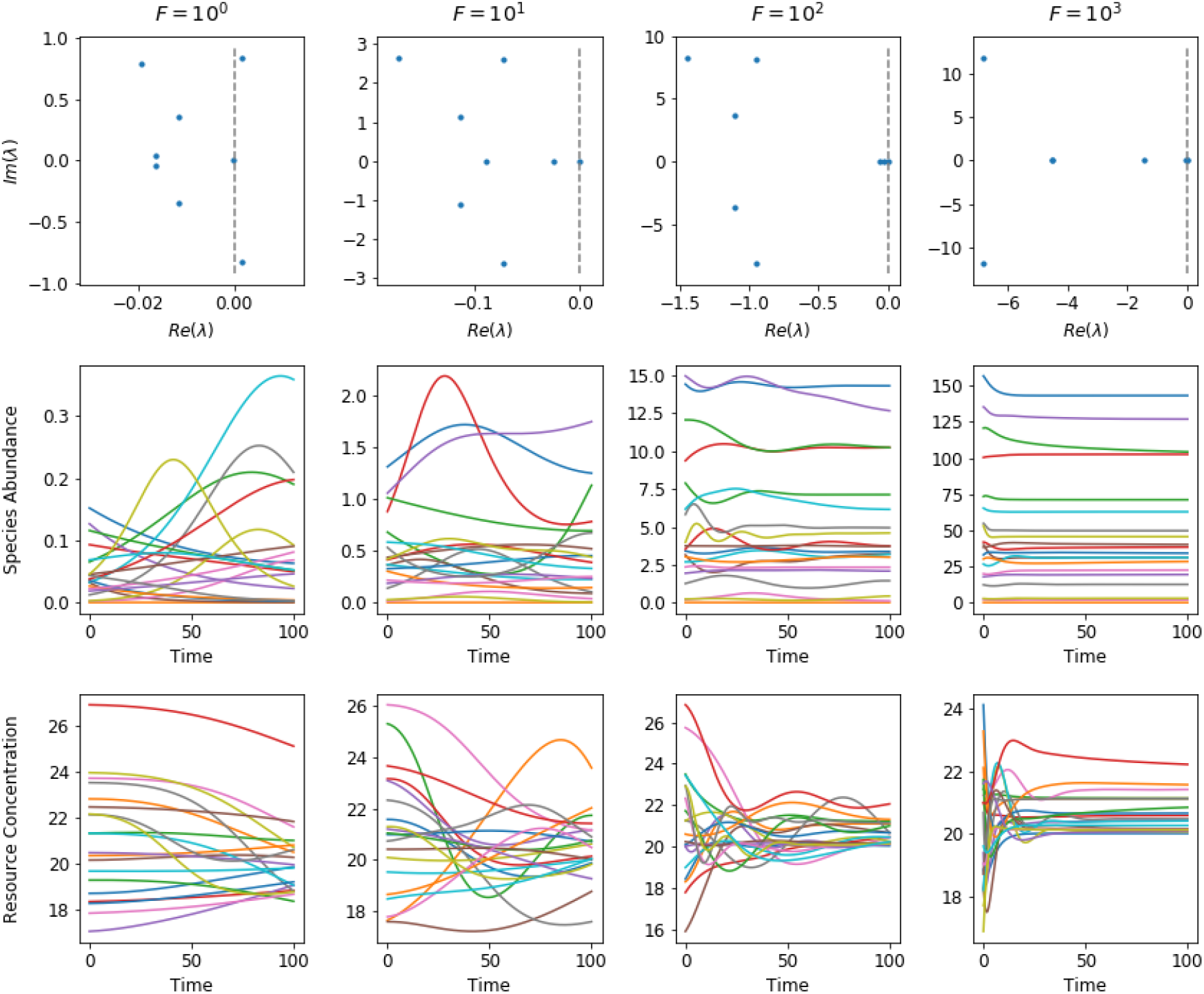
Resource Supply and the Onset of Instability for Substitutable Resources. This example shows four communities of 20 consumer species, each able to use two of 20 distinct resources. Consumption rates are such that the matrix *C* has diagonal entries non-zero, and also one other entry non-zero for each row and column. Entries are again drawn from absolute values of normal distributions.Inflow and mortality rates are chosen so that equilibrium concentrations and abundances are also randomly drawn from the absolute values of a normal distribution, and the exchange matrix Θ has a connectance =0.2, with non-zero entries drawn from the absolute values of a Gaussian with mean zero and sd one. Resource supply rates, 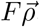, are increased by a factor of ten from each panel to the next, reflected in the increase of consumer abundances by the same factor. The top panels illustrate the same phenomenon by focusing on the eigenvalue spectrum of the Jacobian near the equilibrium in each case. As resource supply is increased, the spectrum loses eigenvalues with positive real part.

**Figure 3:**
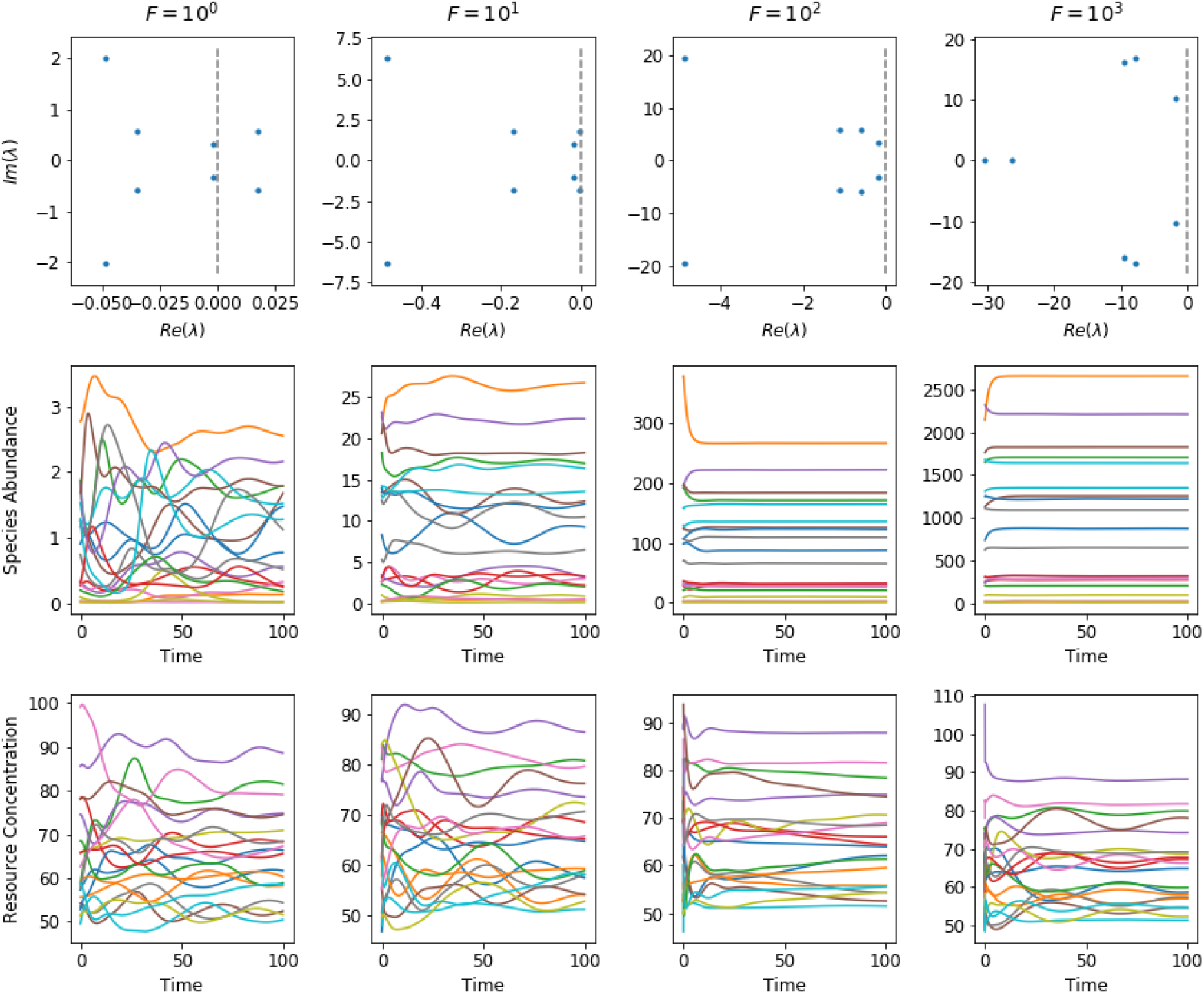
Resource Supply and the Onset of Instability for Essential Resources. This example shows four communties of 20 consumer species, each using an essential combination of multiple resources drawn from a set of 20 distinct resources. The matrix *K*_*αi*_ describing how many units of resource *α* are needed by species *i* was chosen to have a connectance of 0.3, and each entry is randomly drawn from a list containing 1, 2 and 3 with different weightings. I.e. each species uses on average 6 resources, and requires either one, two or three units of each, with one unit being the most likely in this example, followed by two and three. From left to right we show the effect of changing resource supply on the dynamics of this community near equilibrium. Resource supply rates, 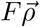, are increased by a factor of ten from each panel to the next, reflected in the increase of consumer abundances by the same factor. The top panels illustrate the same phenomenon by focusing on the eigenvalue spectrum of the Jacobian near the equilibrium in each case. As resource supply is increased, the spectrum loses eigenvalues with positive real part.

#### 3.1.1 Specialism

We now explore these criteria and their intepretation in each of our three example models. The specialist consumer model rewritten using matrix-vector notation is

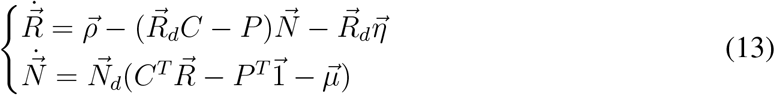

Where *C* is a diagonal matrix.

The three conditions for stability become

1. *C* is nonsingular
2. 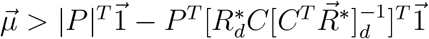
3. (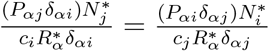 for all *i* and *j*

By definition, this model satisfies the first condition. The second condition holds because *P* is non-negative, 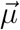 is positive and 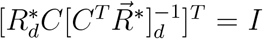. Only the third condition (reciprocity) may or may not hold. If we consider the special case of *c*_*i*_ = *c* and 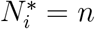 for all *i*, and 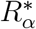 = *r* for all *α*. Then if *P* is symmetric, the third condition will hold, recapitulating the earlier result in (*8*).

#### 3.1.2 Substitutable Resources

The model for substitutable resources rewritten using matrix-vector notation is

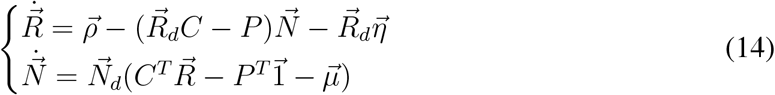

and the sufficient conditions take the form

1. *C* is nonsingular
2. 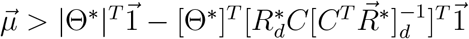
3. 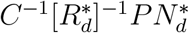 is symmetric

Where 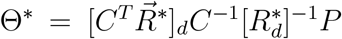. We assume the first condition for our model. This model satisfies the second condition for positive Θ^*\**^ because 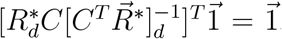. In order to ensure stability, the third condition must be verified using the equilibrium values for a given model.

#### 3.1.3 Colimitation by Multiple Resources

The model in matrix-vector notation is

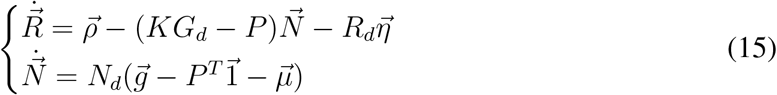

The sufficient conditions for stability become

1. 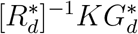 is nonsingular
2. 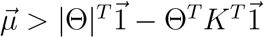
3. 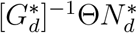 is symmetric

Where Θ = *K*^−1^*P* and *G*_*d*_ is the diagonal matrix with 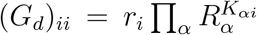. The first condition holds for full-rank *K* and 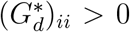, which we assume for our model. The second condition holds for Θ with non-negative entries by noticing that *K* has integer entries is assumed nonsingular so that 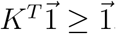. Only reciprocity, the third condition, may fail for this model.

### 3.2 Abundant Resource Inflow Implies Stability

This connection between reciprocity and stability may seem like it must be highly finely tuned—i.e. a very unlikely special case. We now think about the implications for non-reciprocal exchange, and show that (making some additional assumptions) non-reciprocal modes of exchange *can* lead to stable equilibria, but in some cases only when resource inflows are large. Viewed in this light, even being near to a condition of reciprocity may extend the range of environmental contexts in which a given system will have a stable, positive equilbria. In other words, reciprocity guarantees stability in our models for any set of resource supply rates, but a broader range of modes of exchange will be compatible with stability in a given environmental context defined by a finite set of supply rates.

To explore this, we first consider our simplest example above, where each species specializes on a single resource. We choose *η*_*α*_ = 0 and choose *ρ*_*α*_ and *µ*_*i*_ such that each resource has the same equilibrium concentration, 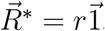, and each species has the same equilibrium abundance 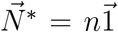. Hence also all consumers have the same growth rate at this equilibrium 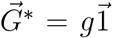, balanced by mortality.

For this model, the condition defining the equilibrium solution is:

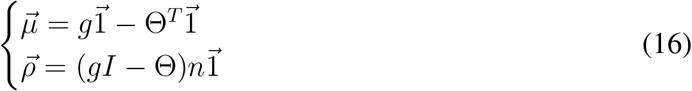

and the Jacobian matrix for small perturbations around this feasible equilibrium solution is:

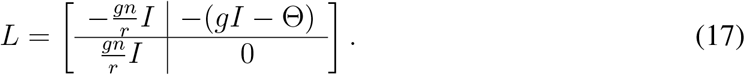

From the block form of this matrix, if *λ* is an eigenvalue of *L*, and *γ* is an eigenvalue of Θ, then these eigenvalues are related by:

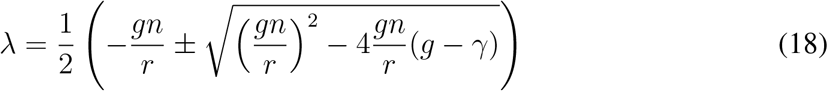

The condition for the equilibrium to be stable to local perturbations (*Re*(*λ*) *<* 0) leads to the following necessary and sufficient condition for all *γ* (*8*):

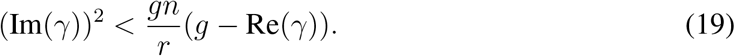

Note that these systems with two species and two resources are always stable. This is because the entries of Θ are all positive, and thus *γ* is real. This fact underlines the importance of considering systems with more than just a few interacting species (*26, 28*).

We now consider what happens when environmental conditions cause the resource influx rate to be reduced. We assume the resource influx is rescaled to 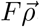, where *F* is a positive real number, and *F* may vary. Then the conditions defining the equilibrium solution become:

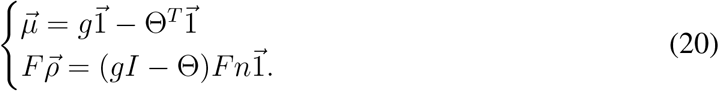

So now the equilibrium abundances of all consumers are *Fn*, and the necessary and sufficient criterion for stability becomes

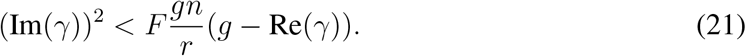

In the limit of scarce resources, i.e. as *F →* 0, the only Θ that will satisfy this inequality are such that Im(*γ*) = 0 for all *γ*, thus restricting the form of Θ to a relatively specific set of matrices, including reciprocal exchange, alongside (for example) exchange networks without feedback loops. On the other hand, when resources are plentiful, i.e. when 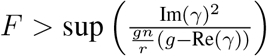, where the supremum is taken over all eigenvalues of Θ, then this system will be stable.

In Appendix C we extend this result, and show that for our more general consumption process positive equilibria will be stable for sufficiently high resource inflow rates, even if they are unstable for lower resource rates. To prove this we state some specific conditions on growth and exchange, but importantly we do not assume reciprocity or any specific structure for the matrix of mutualistic interactions, Θ. To illustrate the implications of this result, we also picked out some explicit examples and solved for their dynamics numerically, shown in Figs 1–3. In these examples, because there is no resource outflow, the feasibility of the equilibrium does not change. Also, we prove in the Appendices that zero can not be an eigenvalue of the Jacobian for our models and so as resource inflow increases, we see a Hopf bifurcation where a stable limit cycle collapses to form a stable equilibrium point. We note that our result contrasts with the well-known paradox of enrichment (*42*), and related results on the effect of spatial subsidies on stability (*32, 41*). We also note that there is no guarantee in our models that the magnitude of the largest eigenvalue of the appropriate Jacobian decreases monotonically with increasing resource inflow, and so it is possible that in some cases there may be phenomena of ‘oscillating’ stability and instability as a function of resource inflow, although always (eventually) resulting in stable equilibria beyond some threshold level.

#### 3.2.1 Transition from Stability to Instability in a Slowly Changing Environment

The effect of a variable environment on individual species extinction has previously been quantified via the raw, direct effect of environmental parameters on species growth rates (*17, 30, 31*), but there is a more limited understanding of how environmental variability might impact collective properties like community stability, in particular when species can exchange resources as well as consume them. We have been agnostic in this paper with respect to how exchange has evolved over time (*4, 48*). But it is clear that generic modes of exchange could evolve (if adaptive) in the context of high resource supply rates, only to lead to community instability when supplies dry up. We provide an explicit example of this phenomenon for a three-species community with specialized consumers. For a sufficiently high set of resource inflow rates, this system has a stable equilibrium—i.e. this is just a specific example of the general case above. We now consider resource inflows to change, but now in the other direction, so that resource inflow is slowly reduced over time. If inflow rates are rescaled by a factor *F*, as above, and this factor varies slowly enough that the system reaches a new equilibrium for each value of *F*, then this system will become unstable for a sufficiently small *F*. We derive the threshold explicitly for this specific system, though the phenomenon is quite general for specialist consumers and matrices Θ with any complex eigenvalues. This suggests that the condition of stability under a range of environmental contexts may be an important filter during the evolution of cooperation.

We begin with no exchange, i.e. Θ = 0. This system always has feasible equilibria for any positive *ρ* and *µ*, and these equilibria is locally stable. Set 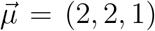, 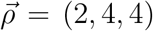. Then 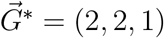, 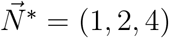 and with 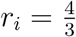 for all *i*, then 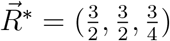.

We next suppose that over time, production is introduced as:

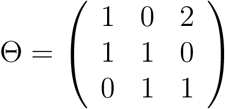

which has eigenvalues, 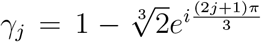 for *j* = 0, 1, 2. The equilibrium values of this system with production, Θ, are 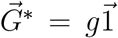 with *g* = 4, 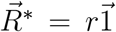 with *r* = 3 and 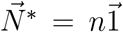 with *n* = 2. This system is stable. The eigenvalues of the Jacobian for this equilibrium are approximately {*−*0.82 *±* 2.86*i, −*1.84 *±* 2.86*i, −*1.33 *±* 1.69*i*} (i.e., the equilibrium is stable to local perturbations).

So we have a system with asymmetric exchange, which nevertheless in this environmental context has a stable equilibrium. We now rescale 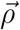 to 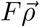, as above so that consumer abundances are rescaled to *n* = 2*F*. The criterion above then tells us that:

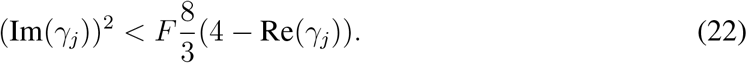

for all *γ*_*i*_. In other words, there is a threshold value of *F* at which this equilibrium will become unstable.

We now define *F*_0_ as this threshold value of *F*, below which (22) fails for some eigenvalue, *γ*_*j*_, of Θ. The largest *F*_0_ comes from the stability criterion applied to *γ*_2_

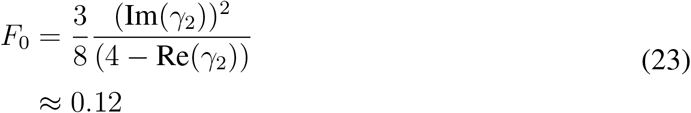

Therefore, for resource inflow levels below *ρ* = *F*_0_ *×* (2, 4, 4), the system with production defined by Θ is unstable. However, the original system with no resource exchange, where Θ = 0, would still have had a positive, stable equilibrium under these reduced resource inflow conditions, implying that (with the benefit of hindsight) the addition of exchange was detrimental to stability.

## 4 Discussion

Our study has identified two important connections between resource exchange and community stability, in the context of consumption of a set of essential resources. Our approach builds on classic consumer-resource dynamics (*1, 11, 49–51*), but incorporates the possibility of arbitrary exchanges of resources between consumers, which could be thought of in terms of the recycling of biomass into usable matter following mortality. Using this framework, we first generalized an earlier connection between pairwise reciprocity in the exchange of resources and community stability for a complex ecological system (*8*). Specifically, reciprocity of resource exchange is a sufficient condition for the local stability of a positive equilibrium, even when consumption requirements of species are a complicated combination of distinct resource types. When reciprocity is absent in our models, the instability that may occur is not the result of differences between impact and consumption vectors (*27, 51*), but due to the exchange process. As long as reciprocity holds, the strength of interactions and dependencies between species can be arbitrarily strong. The difference between this conclusion and the classic results arising from pairwise interactions reinforces the importance of considering explicit mechanisms underlying species interactions—even to gain an understanding of quite basic properties like the local stability of equilibria.

Our second result identifies a connection between the environmental context, characterized by the rates of resource supply from outside, and community stability. We proved that generic modes of exchange lead to stable equilibria if resource supply rates are sufficiently high. When resources are scarce, many modes of exchange will lead to instability, but in times of sufficient resource supply any mode of exchange can lead to a stable equilibrium, in contrast to classic work based on smaller, but trophically-structured networks (*42*). This well-known paradox of enrichment considers the interaction of nutrients, prey, and predators, and for a certain set of functional forms for predator-prey interactions we know that increasing nutrient availability for the prey species will lead to instability. Our models do not not have this three-level trophic interaction, but do allow for a much more complex ‘horizontal’ network of exchange. And yet we find, for essentially any form of this horizontal structure, that a large enough increase in external resource inflow will eventually lead to stability.

We know that models using pairwise interactions are not, as no models are, flawless representations of the real world. To enhance the faithfulness of the representation of reality, it is reasonable to consider incorporating mechanism in a model. The results from our consumer-resource model that contradict those found using models of pairwise interactions, do not imply that such models are invalid, but rather, our model, with its distinct characteristics, provides an important alternative view.

Taken in combination, our two results suggest some extensions to the dictionary of how to be stable: either have a special network structure for positive interactions, or find a habitat with large resource inflows. Clearly there is also an interplay between these results—it may be that even being close to a special network structure, like reciprocal exchange, even if not being exactly at that point, will allow for coexistence across a wider range of resource inflows than a more generic network of exchange. These results also bear comparison with the classic results for pairwise interactions. An intuitive reason for the instability in systems with direct species interactions described by May and others (*2, 37*) is that if there are sufficiently many randomly-placed links between species, and they are strong enough, then these interactions will tend to generate destabilizing feedbacks. It should be noted that an unrealistic type of instability resulting from unbounded growth can be seen in models representing mutualism as direct species interactions. Avoiding feedbacks therefore is a reasonable lesson to take from these systems. Some of the intuition behind these results is similar—for example in the case of species that specialize on a single resource, acyclic exchange (like reciprocity) is an example of the special modes of exchange that will guarantee stability under all environmental contexts. On the other hand, the idea that reciprocity leads to stability is somewhat invisible in the direct interactions approach, where the received wisdom has been that asymmetric mutualisms are more likely to lead to stability (*6*). Yet here we have shown that reciprocity is a general way to ensure stability across different modes of resource consumption.

Finally, we did not make reference to the evolution of cooperation, which in this context would allow for changes in production and exchange over time (*4*), and has been used to shed light on mutalistic ecological interactions among multiple species (*55*). In the right circumstances, these evolutionary processes can lead to exact pairwise reciprocity (*3, 48, 55*), though also to other network structures. For example it is possible for evolutionary processes to result in cyclic non-extractive exchange, known as indirect reciprocity (*7*). Our analysis demonstrates that ecological stability may be a non-trivial criterion along the way for any evolving system of cooperating species. For example, cyclic exchange in our models can lead to instability for certain ranges of resource inflow. This isn’t so much inconsistent with those evolutionary results, but adds a new criterion for stable coexistence at a positive equilibrium, explicitly depending on environmental context. Finally, we note that studies of the evolution of mutualistic exchange have invoked “biological markets” (*29, 47*), in analogy with economic markets. We therefore propose that the criteria for ecological stability derived here may also be important, or at least will provoke interesting exploration, in other complex systems where markets structure the form of exchange networks (*14, 23, 25, 40*).

## Acknowledgements

JOD acknowledges the Simons Foundation Grant #376199, McDonnell Foundation Grant #220020439, NSF #DEB1557192.

## Appendices

### A Feasible Equilibrium Solutions

We consider the following equilibrium solutions of our general model, Eq. (12), in the main text:

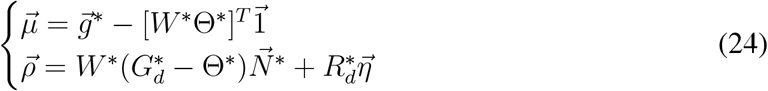

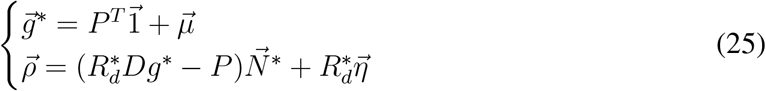

Positive values of *Dg*_*αi*_ ensure the existence of positive 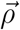 and 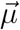. If *Dg*_*αi*_ > 0 for all *α, i*, then there exists 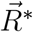 such that 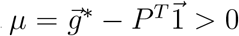 and 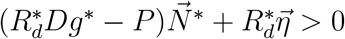. Also, given 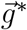, sufficiently small production *P* allows for positive 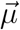 and especially when *Dg*^*^ > 0 or 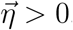.

#### A.1 Specialism

We next consider the equilibrium solutions of one of our specific cases, Eq. (1) in the main text:

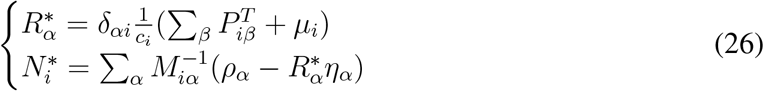

where 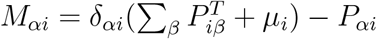.

The equilibrium solutions for 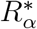 should always be feasible in this model. All of the parameters in the equation for 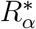 are positive. For 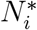, the matrix *M* is an *M-matrix* and so is inverse-positive, meaning that all of the entries of *M*^−1^ are positive. For a given set of 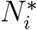, we may find *ρ*_*α*_ and *η*_*α*_ such that 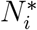 is a solution.

#### A.2 Substitutable Resources

Here we consider the following equilibrium solutions of Eq (3) in the main text:

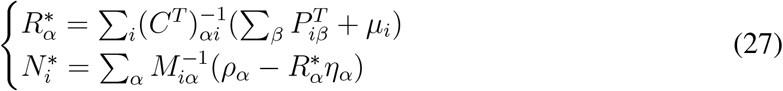

Where 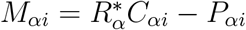.

Although in the model, feasibility is not guaranteed for 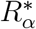, for any given set of positve 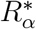, there exists a set of *µ*_*i*_ such that 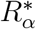 is a solution. For a given set of 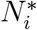, we may find *ρ*_*α*_ and *η*_*α*_ such that 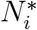 is a solution.

#### A.3 Colimitation by Multiple Resources

We consider the following equilibrium solutions of Eq.(5) in the main text:

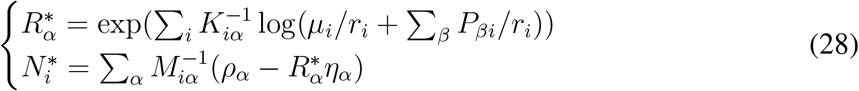

where the matrix *M* is given by *M*_*αi*_ = *K*_*αi*_(*µ*_*i*_ + Σ_*β*_ *P*_*βi*_) − *P*_*αi*_. Clearly not all combinations of *ρ*, *µ*, *r*_*i*_, *K* and *P* will lead to positive (so-called feasible) solutions. On the other hand, all positive values 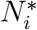 are possible for some choice of resource inflow *ρ*_*α*_. Conversely, not all resource concentrations are possible—the recycling of resources into the common pool typically places lower bounds on equilibrium resource concentrations for a given *K* and *P*.

More explicitly, consider

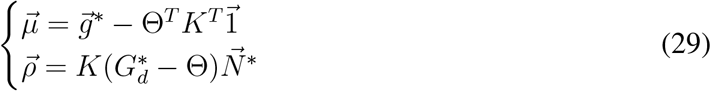

where 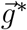 is a vector of equilibrium growth rates 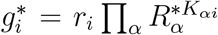, definined in the same way as in the main text, and 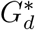 is a diagonal matrix with diagonal elements equal to 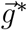.

When production (i.e. the matrix Θ) is non-zero, not all pairs, 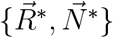, are possible positive (also known as feasible) equillibrium solutions. The primary constraint is that the input rate 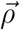 and mortality rate of consumers 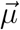 must both be non-negative in order to have a plausible biological interpretation. So this means that the question is how to determine which 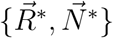 are solutions for positive 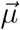 and 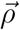, given a fixed set of resource requirements *K* and production defined by the matrix Θ.

First, we note that given a vector 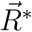, which may or may not be a feasible solution with positive 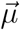, there is always some positive constant, *a* such that 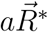 *will* arise from a positive vector 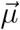. So not all 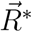 are possible, but there is always a set of resource densities in the same ratios that is a feasible solution. Conversely, all positive 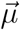 lead to positive 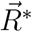.

Second, given 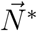, and large enough 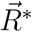, the product 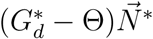 will be positive and thus so will 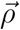. So any positive 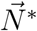 is possible, given a sufficiently large set of resource densities 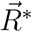. In summary, we can rule out the existence of pathological combinations of *K* and *P* —all pairs allow for at least some range of feasible equilibria.

### B Local Stability for Arbitrary Consumption Requirements

First, writing production in terms of the matrix Θ as defined in Eq. (11) in the main text, the Jacobian at equilibrium is given by

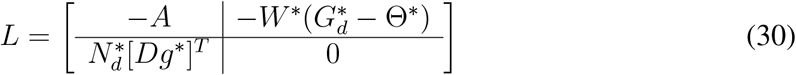

Where *A* is defined as

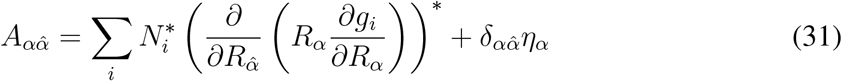

If all the real parts of the eigenvalues of this Jacobian are negative, then this equilibrium is locally stable.

To start with, we show that *λ* = 0 is not an eigenvalue of *L*. Assume it is, then there is an eigenvector,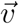, associated with *λ* = 0. Write 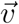 as

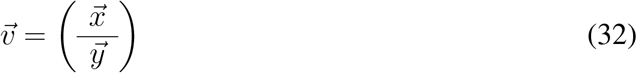

where 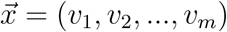 and 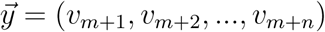 then

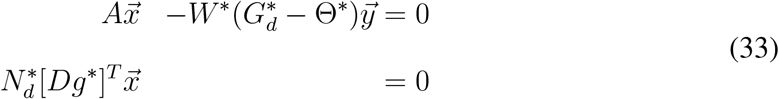

so that 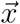 must be in the nullspace of 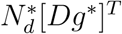. However, for feasible equilibria, 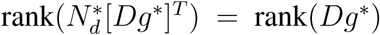. We assume rank(*Dg*^*^) = *m* and so with *Dg*^*^ an *m × m* matrix, then we must have 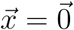. Similarly, 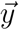 must be in the nullspace of 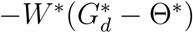 and thus 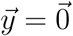 so long as 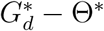 has no zero eigenvalues.

To determine conditions under which the real parts of all the eigenvalues of *L* are negative, we first note that the eigenvalue equation det(*L − λI*) = 0 is given by

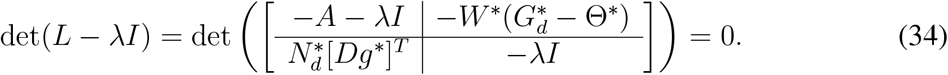

So long as *λ* ≠ 0, then *−λI* is invertible, and so

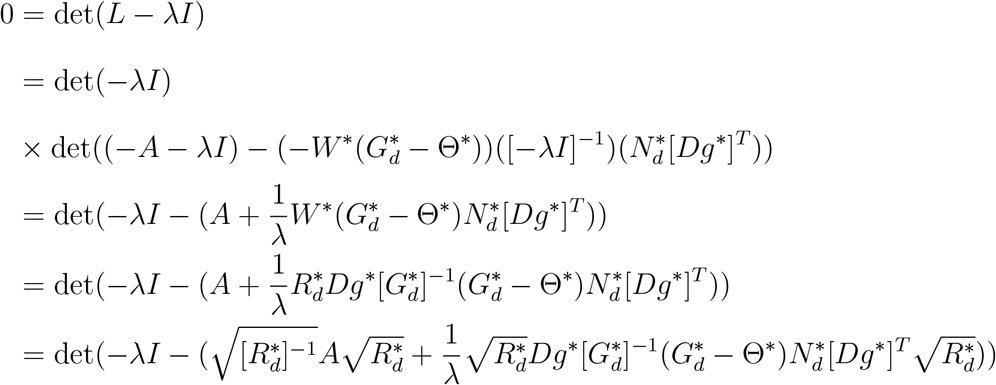

We will next assume *Re*(*λ*) *≥* 0 and look for a contradiction. We will also assume that 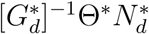 is symmetric, which we interpret in terms of reciprocity between species.

We wish to find a contradiction by proving the eigenvalues of 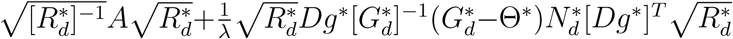 have positive real part and so it will be sufficient to show that the Hermitian part, 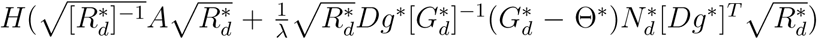, is positive definite. We will do this by looking at the two matrices 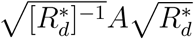 and 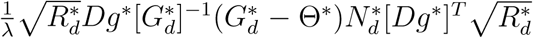 seperately. For 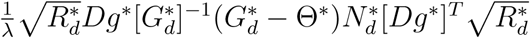

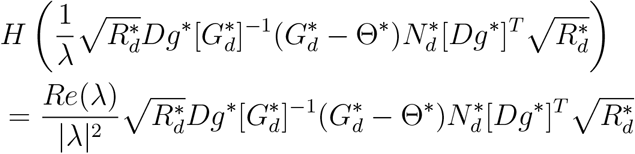

which is ^*T*^ congruent to 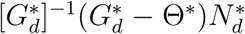 and so we want the eigenvalues of 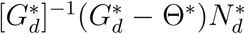 to be all positive. 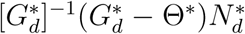 is symmetric, and is ^*T*^ congruent to, and thus has the same inertia as 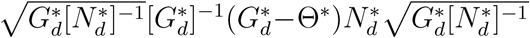 which has the same eigenvalues as 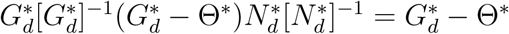.

Thus we wish to find a condition ensuring all eigenvalues of 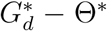 are positive. From Gershgorin’s theorem, this will be true if 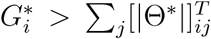 for all *i*. Using that for feasible equilibria, 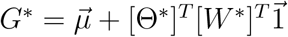, the condition becomes

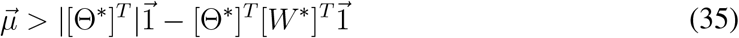

Now looking at the Hermitian part of 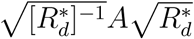. We can write this as 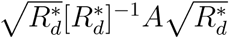 and so if the Hermitian part of 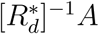 is positive definite so will be 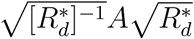. Note that 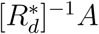 is symmetric.

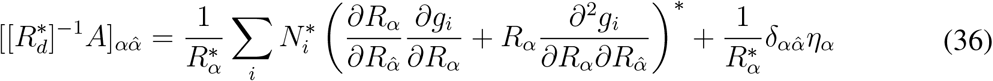

#### B.1 The Example Models Satisfy 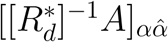 is Positive Definite Specialism

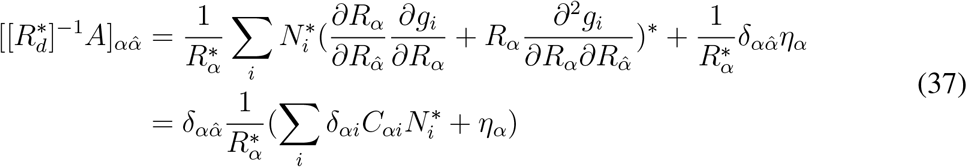

which is a positive definite matrix.

##### Substitutable Resources

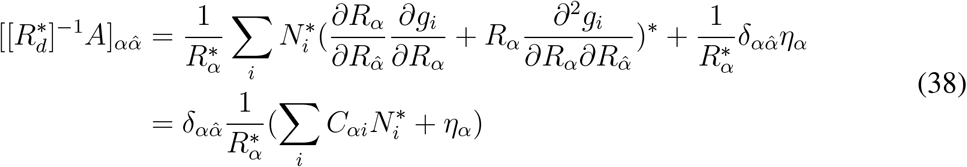

which is a positive definite matrix.

##### Colimitation by Multiple Resources

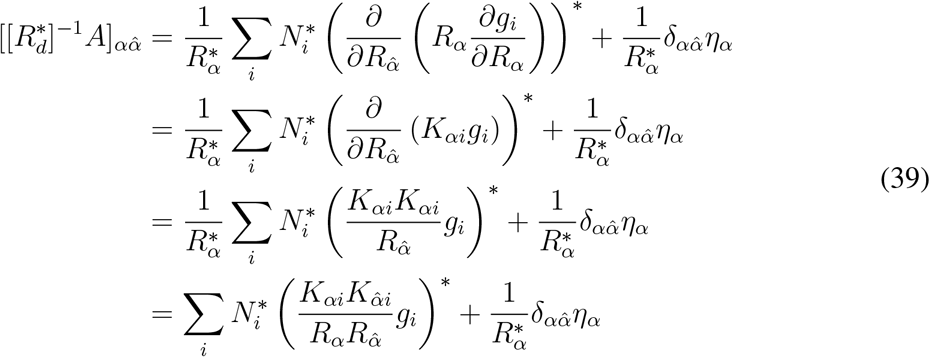

and so

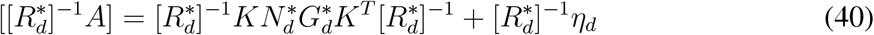

is the sum of two positive definite matrices.

### C Abundant Resource Inflow Implies Stability

#### C.1 Generalized Consumption with No Resource Outflow

Here we consider our model for generalized consumption requirements with no resource outflow, so that 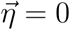. From section B we have the eigenvalue equation

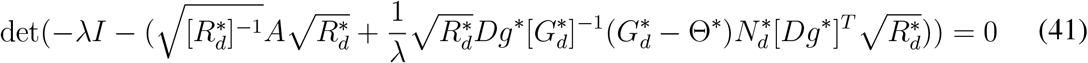

which is equivalent to

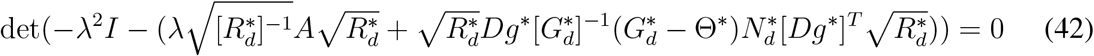

Now consider increasing resource inflow so that 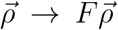 for positive real *F*. Then by Eq. (10) in the main text, the new equilibrium species abundances become 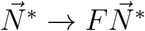 and the eigenvalue equation becomes

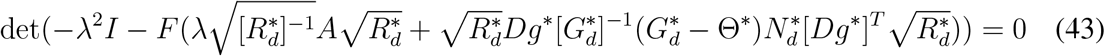

Now for some notation changes. First lets drop the *\** (which from now on will denote complex conjugate transpose). Next define *U* as 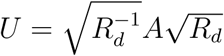 and *V* as 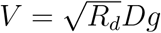 and we rewrite the eigenvalue equation as

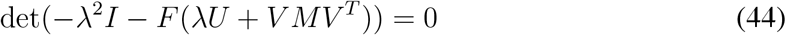

Where 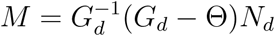.

We wish to show that for large enough *F*, *λ* such that *Re*(*λ*) *≥* 0 cannot be a solution. We will not need to consider which *λ* actually solve (44). We will only need to eliminate the ones with positive real part. We assume, as before, that *V* is nonsingular and that *U* is positive definite. We also assume that the field of values of *M*, 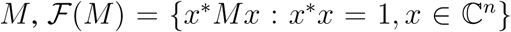 is contained strictly in the right half of the complex plane. This condition is somewhat stricter than that used in conjunction with symmetry for the stability proof in the paper, which was that *M* had all positive eigenvalues.

Let *λ* = *a* + *bi*. Assume *a ≥* 0 and, by symmetry, it is only necessary to consider *b ≥* 0 so assume that as well. We show for such *λ* that there exists an *F*_*λ*_ such that for all *F > F*_*λ*_, 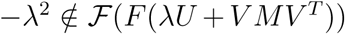 and thus *−λ*^2^ cannot be an eigenvalue of *F* (*λU* + *V MV*^*T*^). Then we argue that we can bound our {*F*_*λ*_}.

Note that from the properties of field of values we have

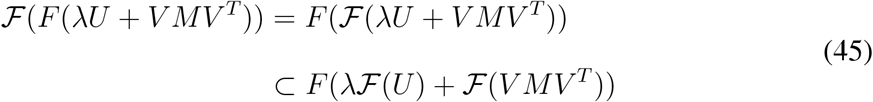

We will essentially show that we can use *F* to push 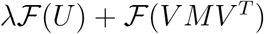 too far from the origin, beyond |−*λ*^2^|. Because of this, it is the values of 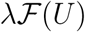 and 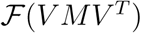 with minimum modulus that will be important.

First, we want to describe 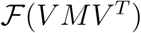.

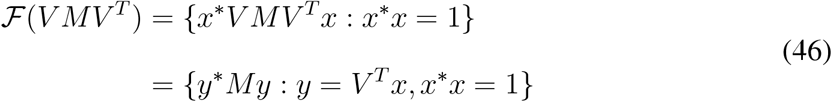

For every *y* = *V*^*T*^*x* with *x*^*^*x* = 1, we have that there exists an 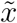 with 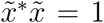 such that 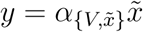 for some real number 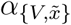. And so continuing

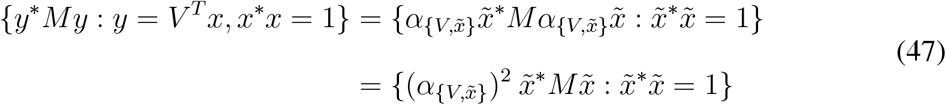

so that every point in the field of values 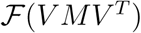 is a point in 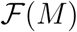 stretched/shrunk by some value 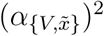. We want to be sure 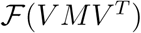 is contained in the right half of the complex plane and so need to consider the shrink. Because *V*^*T*^ is non-singular, from the singular value decomposition of *V*

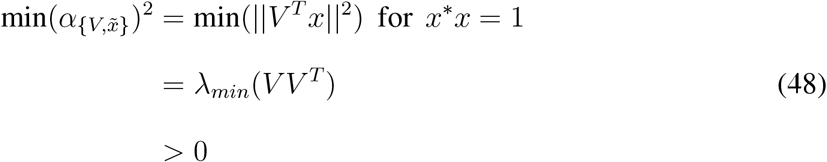

Where *λ*_*min*_(*V V*^*T*^) is the smallest eigenvalue of *VV*^*T*^. Now because 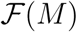 is in the right half of the complex plane and *λ*_*min*_(*V V*^*T*^) *>* 0 we may conclude that 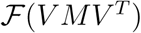 is contained in the right half of the complex plane. Because a field of values is a compact, convex set, which for a real matrix will be symmetric about the real line, we may without loss of generality, assume that 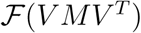 is a closed disk, *D*, centered at *c* with radius *r*. And because we know that every value in 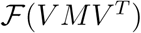 has positive real part, we have that *r < c*.

**Figure 4:**
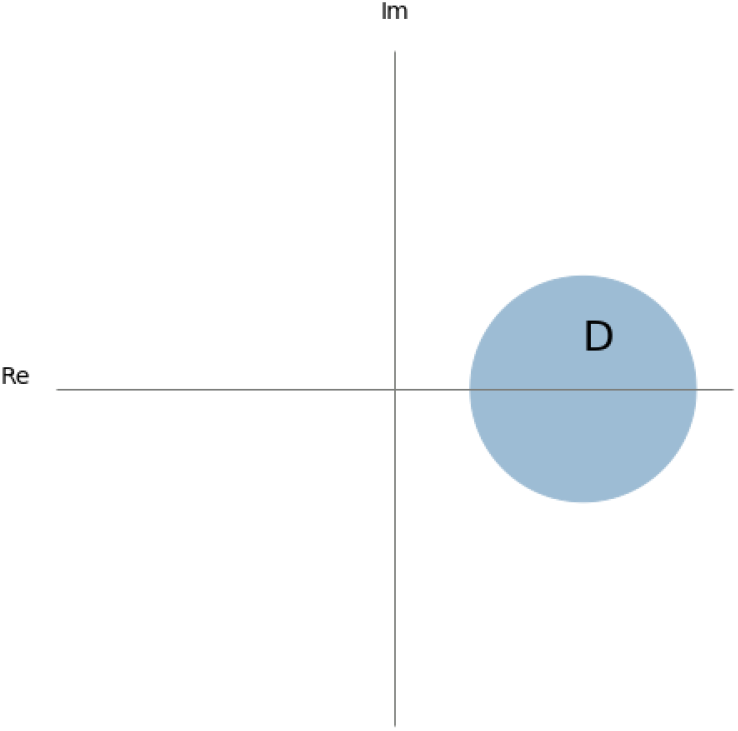
The disk, *D*, which represents 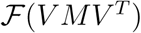. It is symmetric about the real line and is contained strictly in the right half of the complex plane

Now lets look at 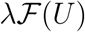. Because *U* is positive definite, 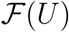 is just a closed interval of positive real numbers, [*κ*_0_, *κ*_1_]. So to summarize, for each *λ* = *a* + *bi* with *a, b ≥* 0 we are looking for an *F*_*λ*_ such that *F* > *F*_*λ*_ implies *−λ*^2^ ∉ *F* ([*κ*_0_*λ, κ*_1_*λ*] + *D*) where we define the sum of two sets by: *S*_1_ + *S*_2_ = {*s*_1_ + *s*_2_: *s*_1_ *∈ S*_1_, *s*_2_ *∈ S*_2_}

The distance of [*κ*_0_*λ, κ*_1_*λ*] + *D* to the origin can be no less than *c − r* and so we can find *F*_*λ*_ such that

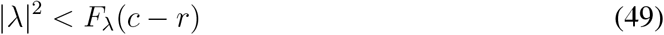

Now we will show that we only need to consider *λ* in a bounded region. First note that for *λ* = *|λ|e*^*iθ*^ such that 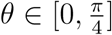, we have 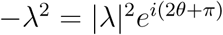 with 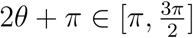 and so for all *F*, *−λ*^2^ *∉ F* ([*κ*_0_*λ, κ*_1_*λ*] + *D*).

Consider, *λ* = *a* + *bi* such that *κ*_0_*b > r*, then [*κ*_0_*λ, κ*_1_*λ*] + *D* lies entirely above the real axis, however, for all *λ* = *a* + *bi* such that *a, b ≥* 0, *−λ*^2^ is in the lower half of the complex plane. Thus we have a region, *R*, bounded by the imaginary axis, the ray 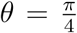 and the line 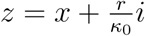.

**Figure 5:**
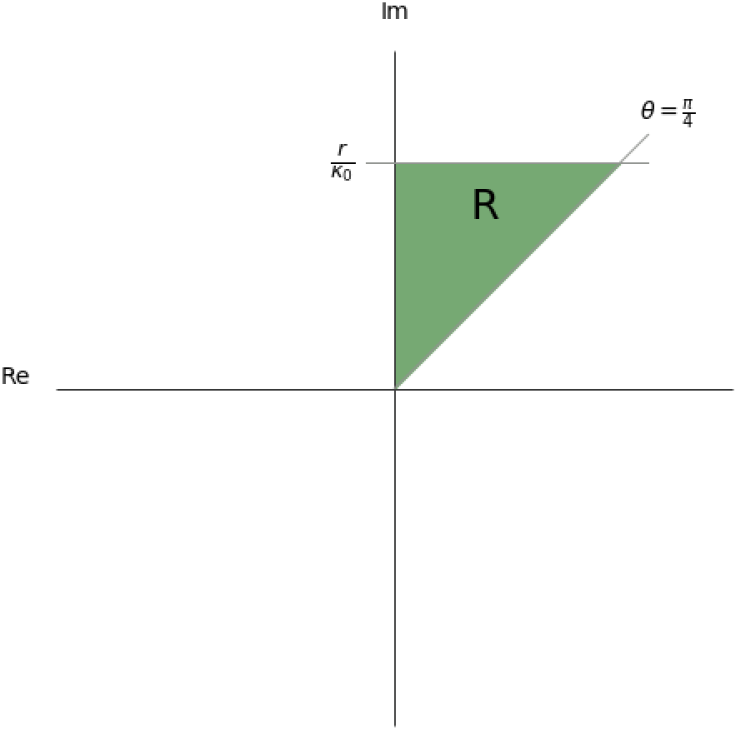
The region, *R*, in the complex plane. For *λ* outside of this region, −*λ*^2^ *∉ F* ([*κ*_0_*λ, κ*_1_*λ*] + *D*) for all *F >* 0

From (63), larger magnitude *λ* require larger *F*_*λ*_. Consider *λ*_0_ = *a* + *bi* such that *a* = *b* and 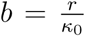. Then *|λ*_0_| > |*λ|* for any *λ* in *R* and so can be used along with (63) to give an upper bound, *F*_0_, for {*F*_*λ*_}

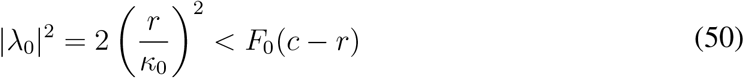

And so 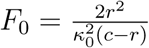 is the desired bound for {*F*_*λ*_}.

#### C.2 Generalized Consumption with Resource Outflow

Here we consider our model for generalized consumption requirements with resource outflow, so that 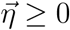. From above, we have the eigenvalue equation

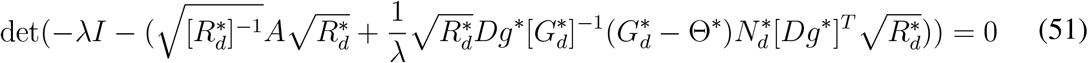

From here on, although all values will continue to be equilibrium values, we drop the ^*\**^ notation, which will from now on be used to indicate complex conjugate transpose. The eigenvalue equation is equivalent to

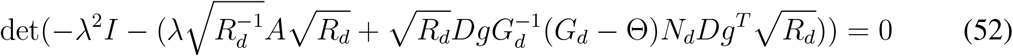

For this proof, we will need to alter some of the previous assumptions. The first involves the assumption that

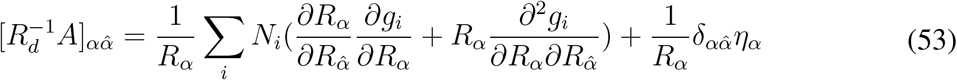

is a positive definite matrix. Our altered assumption is that

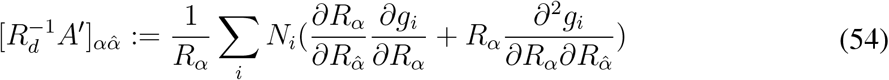

is positive definite. Note that this assumption implies the previous one. All of our example models satisfy this strengthened assumption. Define 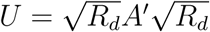 and 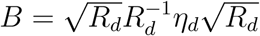. Also define 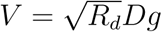. The eigenvalue equation now can be written

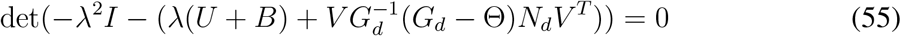

Now consider increasing resource inflow so that 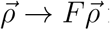 for positive real *F*

Define 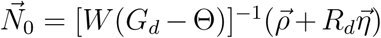 and 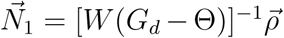 where 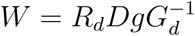 as in the main text. Then the new equilibrium species abundances become

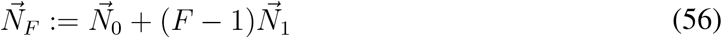

Finally define 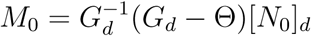 and 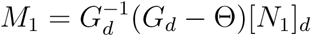 so that

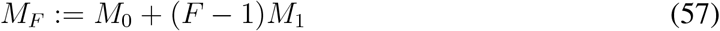

and the new eigenvalue equation is

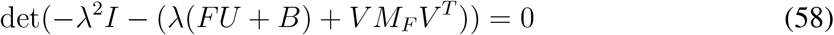

We wish to show that for large enough *F*, *λ* such that *Re*(*λ*) *≥* 0 cannot be a solution. We will not need to consider which *λ* actually solve (58). We will only need to eliminate the ones with positive real part. We assume, as before, that *V* is nonsingular. We also assume that the field of values of *M*_1_, 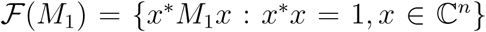 is contained strictly in the right half of the complex plane. Note that in the absence of outflow, 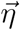, this assumption is equivalent to the one used in the stability proof for the 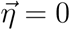 case, where it was assumed that 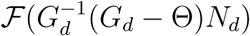 is contained strictly in the right half of the complex plane.

Let *λ* = *x* + *yi*. Assume *x ≥* 0 and, by symmetry, it is only necessary to consider *y ≥* 0, so assume that as well. We show for such *λ* that there exists an *F*_*λ*_ such that for all *F > F*_*λ*_

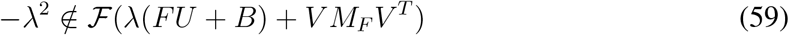

and thus *−λ*^2^ cannot be an eigenvalue of

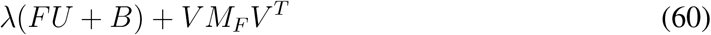

Then we argue that we can bound our {*F*_*λ*_}.

Note that from the properties of field of values we have

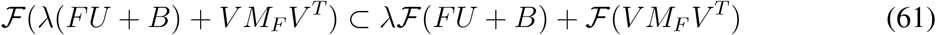

where we define the sum of two sets by: *S*_1_ + *S*_2_ = {*s*_1_ + *s*_2_: *s*_1_ *∈ S*_1_, *s*_2_ *∈ S*_2_}.

We will essentially show that we can use *F* to push 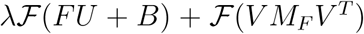 too far from the origin, beyond *|− λ*^2^|.

We wish for the elements of 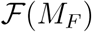 to have positive real parts. Now because 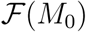 is bounded, 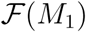 is in the right half of the complex plane, and field of values is subadditive, there exists an *F*_*ρ*_ such that 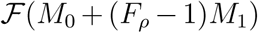 is in the right half of the complex plane. Now

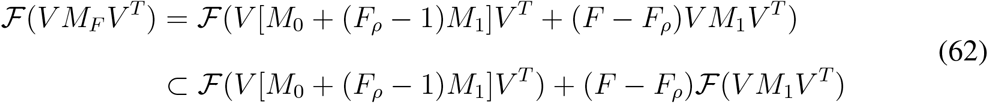

From the argument given for the proof of the 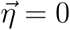 case, because *V* is nonsingular, we may conclude that both 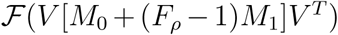 and 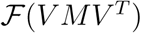 are contained in the right half of the complex plane. Because a field of values is a compact, convex set, which for a real matrix will be symmetric about the real line, we may without loss of generality, assume that 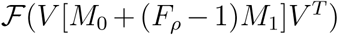 and 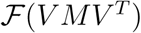 are closed disks, *D*_1_ and *D*_2_, with centers at *c*_1_ and *c*_2_, radii *r*_1_ and *r*_2_ respectively. And because we know that they are in the right half of the complex plane, we have that *r*_1_ *< c*_1_ and *r*_2_ *< c*_2_.

Now lets look at 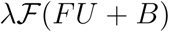. Because *U* and *B* are positive definite, the field of values of each is just a closed interval of positive real numbers, [*u*_0_, *u*_1_] and [*b*_0_, *b*_1_] respectively. So to summarize, for each *λ* = *x* + *yi* with *x, y ≥* 0 we are looking for an *F*_*λ*_ such that *F* > *F*_*λ*_ > *F*_*ρ*_ implies *−λ*^2^ *∉ λ*[*Fu*_0_ + *b*_0_, *Fu*_1_ + *b*_1_] + *D*_1_ + (*F* − *F*_*ρ*_)*D*_2_.

Because all the elements of *λ*[*Fu*_0_ + *b*_0_, *Fu*_1_ + *b*_1_] + *D*_1_ have positive real parts, the distance of *λ*[*Fu*_0_ + *b*_0_, *Fu*_1_ + *b*_1_] + *D*_1_ + (*F* − *F*_*ρ*_)*D*_2_ to the origin can be no less than the distance of (*F* − *F*_*ρ*_)*D*_2_ to the origin, (*F* − *F*_*ρ*_)(*c*_2_ *− r*_2_). So we can find *F*_*λ*_ such that

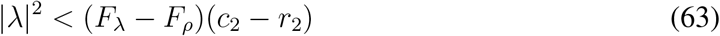

Now we will show that we only need to consider *λ* in a bounded region. First note that for *λ* = *|λ|e*^*iθ*^ such that 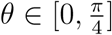, we have *−λ*^2^ = *|λ|*^2^*e*^*i*(2*θ*+*π*)^ with 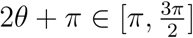 and so for all *F*, *−λ*^2^ *∉ λ*[*Fu*_0_ + *b*_0_, *Fu*_1_ + *b*_1_] + *D*_1_ + (*F* − *F*_*ρ*_)*D*_2_.

Consider, *λ* = *x* + *yi* such that *y > κ* where 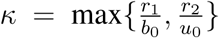 then (*Fu*_0_ + *b*_0_)*y > r*_1_ + (*F* − *F*_*ρ*_)*r*_2_, and *λ*[*Fu*_0_ + *b*_0_, *Fu*_1_ + *b*_1_] + *D*_1_ + (*F* − *F*_*ρ*_)*D*_2_ lies entirely above the real axis for all *F* > *F*_*ρ*_, however, for all *λ* = *x* + *yi* such that *x, y ≥* 0, *−λ*^2^ is in the lower half of the complex plane. Thus we have a region, *R*, bounded by the imaginary axis, the ray 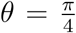 and the line *z* = *x* + *κi*.

**Figure 6:**
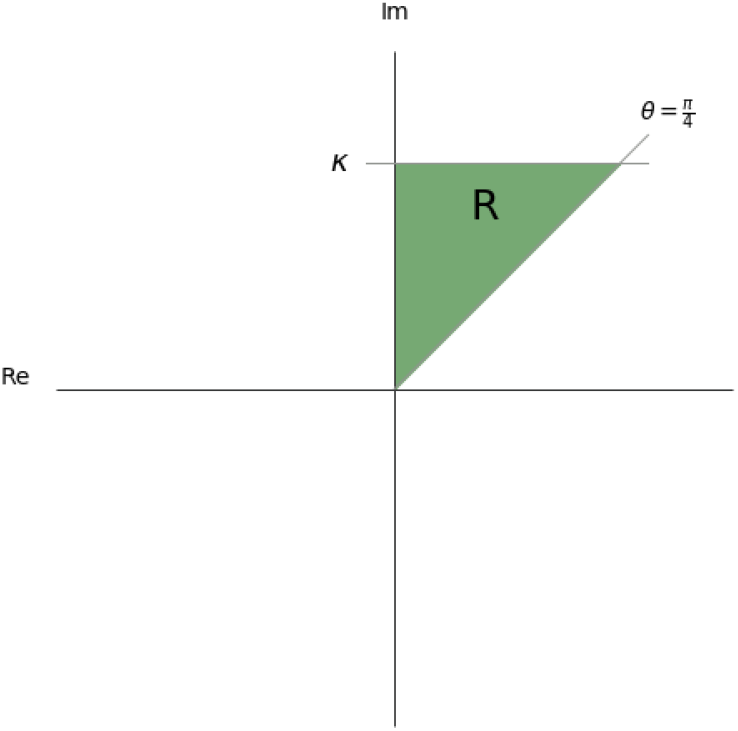
The region, *R*, in the complex plane. For *λ* outside of this region, *−λ*^2^ *∉ λ*[*Fu*_0_ + *b*_0_, *Fu*_1_ + *b*_1_] + *D*_1_ + (*F* − *F*_*ρ*_)*D*_2_ for all *F* > *F*_*ρ*_

From (63), larger magnitude *λ* require larger *F*_*λ*_. Consider *λ*_0_ = *x* + *yi* such that *x* = *y* and *y* = *κ*. Then *|λ*_0_*| > |λ|* for any *λ* in *R* and so can be used along with (63) to give an upper bound, *F*_0_, for {*F*_*λ*_}

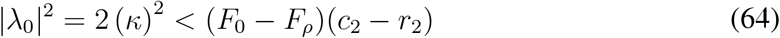

And so 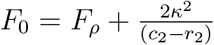 is the desired bound for {*F*_*λ*_}.

### D Local Stability when *n* > *m*

For this section, we assume that the number of resources is greater than the number of consumer species. As in Section B, the Jacobian at equilibrium is given by

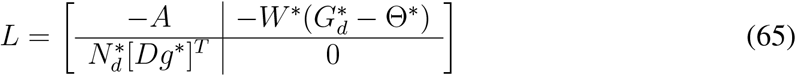

Where *A* is defined as

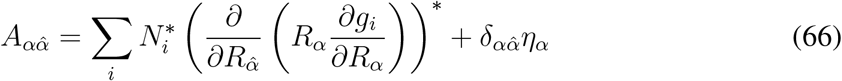

If all the real parts of the eigenvalues of this Jacobian are negative, then this equilibrium is locally stable.

To start with, we show that *λ* = 0 is not an eigenvalue of *L*. We have the eigenvalue equation

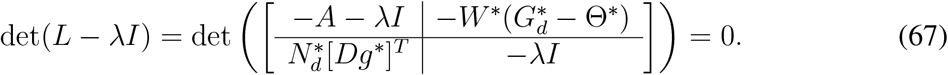

Then, when *λ* is not an eigenvalue of *−A*

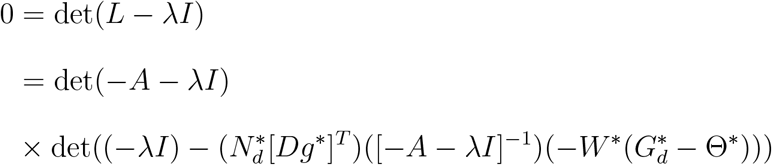

Now, we assume *λ* = 0 is an eigenvalue of *L* and we find a contradiction. Because we assume that *−A* is positive definite, we know *λ* = 0 is not an eigenvalue of *−A*, and so using the preceeding equation with *λ* = 0 we write

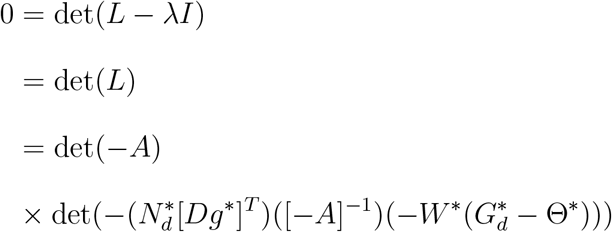

and we have

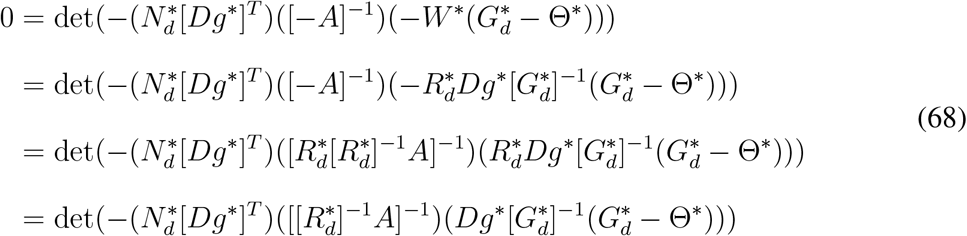

We will now do a rank analysis to show the preceeding equation is false. First, 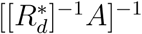 is the inverse of an *n × n*, positive definite matrix and so is also *n × n*, positive definite. Next, note that *Dg*^*^ is an *n × m* matrix of rank *m* and so 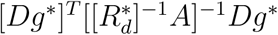 is an *m × m* positive definite matrix. We also have that 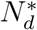 and 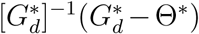 are both *m × m* matrices of rank *m*. And so, taken together these facts imply that the matrix inside the determinant in the preceeding equation is *m × m* with rank *m*, and so the equation must be false.

Now, it remains to show that the real parts of all the eigenvalues of *L* are negative, we first note that the eigenvalue equation det(*L − λI*) = 0 is given by

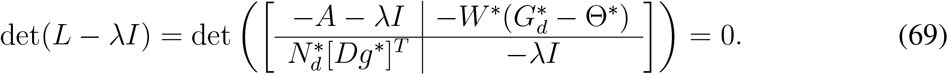

So long as *λ* ≠ 0, then *−λI* is invertible, and so

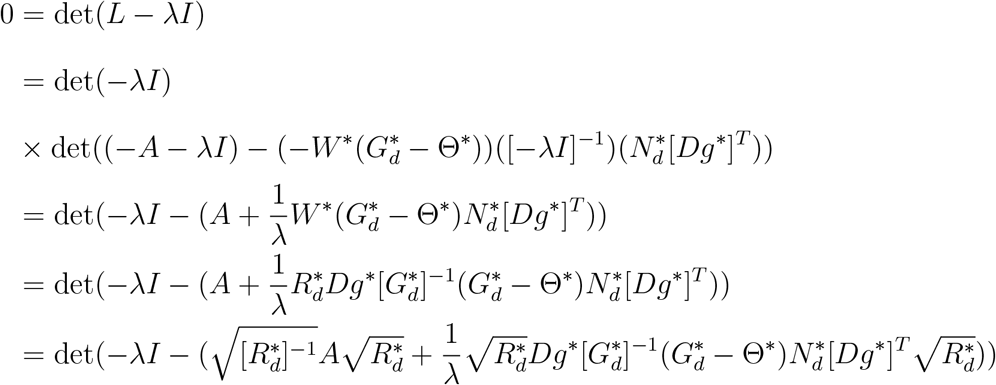

We will next assume *Re*(*λ*) *≥* 0 and look for a contradiction. We will also assume that 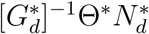 is symmetric, which we interpret in terms of reciprocity between species.

We wish to find a contradiction by proving the eigenvalues of 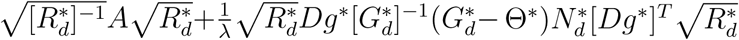 have positive real part and so it will be sufficient to show that the Hermitian part, 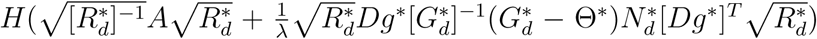, is positive definite. We will do this by looking at the two matrices 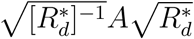 and 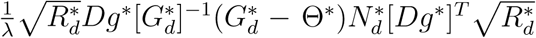 seperately. For 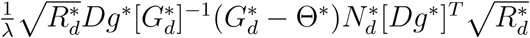

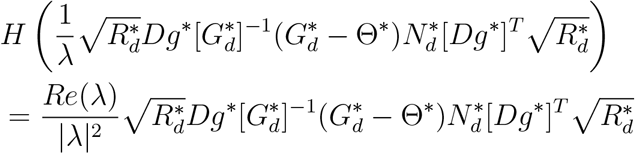

We wish for all the eigenvalues of 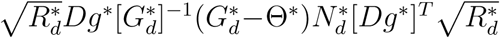 to be non-negative. This will happen if the eigenvalues of 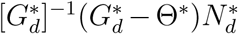 are all positive. 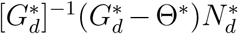 is symmetric, and is ^*T*^ congruent to, and thus has the same inertia as 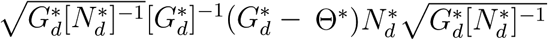 which has the same eigenvalues as 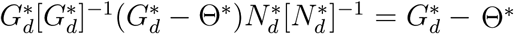.

Thus we wish to find a condition ensuring all eigenvalues of 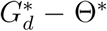 are positive. From Gershgorin’s theorem, this will be true if 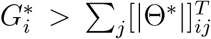 for all *i*. Using that for feasible equilibria, 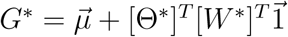, the condition becomes

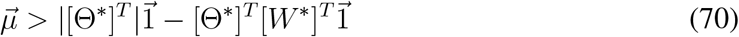

Now looking at the Hermitian part of 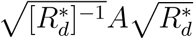. We can write this as 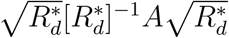 and so if the Hermitian part of 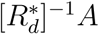 is positive definite so will be 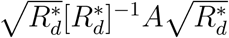. Note that 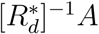 is symmetric.

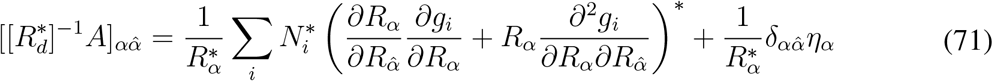

